# Niche and local geography shape the pangenome of wastewater- and livestock-associated *Enterobacteriaceae*

**DOI:** 10.1101/2020.07.23.215756

**Authors:** Liam P. Shaw, Kevin K. Chau, James Kavanagh, Manal AbuOun, Emma Stubberfield, H. Soon Gweon, Leanne Barker, Gillian Rodger, Mike J. Bowes, Alasdair T. M. Hubbard, Hayleah Pickford, Jeremy Swann, Daniel Gilson, Richard P. Smith, Sarah J. Hoosdally, Robert Sebra, Howard Brett, Tim E. A. Peto, Mark J. Bailey, Derrick W. Crook, Daniel S. Read, Muna F. Anjum, A. Sarah Walker, Nicole Stoesser, on behalf of the REHAB consortium

**Affiliations:** Nuffield Department of Medicine, John Radcliffe Hospital, University of Oxford, Oxford, OX3 9DU, UK; Department of Bacteriology, The Animal and Plant Health Agency (APHA), Woodham Lane, Addlestone, Surrey, KT15 3NB, UK; UK Centre for Ecology & Hydrology (UKCEH), Benson Lane, Crowmarsh Gifford, Wallingford, OX10 8BB, UK; School of Biological Sciences, University of Reading, RG6 6AS, UK; NIHR Oxford Biomedical Research Centre; Department of Tropical Disease Biology, Liverpool School of Tropical Medicine, Pembroke Place, Liverpool, L3 5QA, UK; NIHR Health Protection Research Unit in Healthcare Associated Infections and Antimicrobial Resistance at University of Oxford in partnership with Public Health England, Oxford, OX4 9DU, UK; Department of Epidemiological Sciences, The Animal and Plant Health Agency (APHA), Woodham Lane, Addlestone, Surrey, KT15 3NB, UK; Department of Genetics and Genomic Sciences, Icahn School of Medicine at Mount Sinai, New York, NY, 10029, USA; Thames Water Utilities, Clearwater Court, Vastern Road, Reading, RG1 8DB, UK

**Keywords:** antimicrobial resistance (AMR), bacterial genomics, *Enterobacteriaceae*, *Escherichia coli*, pangenomes, environmental bacteria

## Abstract

*Escherichia coli* and other *Enterobacteriaceae* are highly diverse species with ‘open’ pangenomes^1,2^, where genes move intra- and inter-species via horizontal gene transfer^3^. These species can cause clinical infections^4,5^ as well as persist environmentally^6,7^. Environmental populations have been suggested as important reservoirs of antimicrobial resistance (AMR) genes. However, as most analyses focus on clinical isolates^8,9^, the pangenome dynamics of natural populations remain understudied, particularly the role of plasmids. Here, we reconstructed near-complete genomes for 828 *Enterobacteriaceae*, including 553 *Escherichia* spp. and 275 non-*Escherichia* species with 2,293 circularised plasmids in total, collected from nineteen locations (livestock farms and wastewater treatment works in the United Kingdom) within a 30km radius at three timepoints over the course of a year. We find different dynamics for the chromosomal and plasmid-borne components of the pangenome, showing that plasmids have a higher burden of both AMR genes and insertion sequences, and AMR plasmids show evidence of being under stronger selective pressure. Focusing on *E. coli*, we observe that plasmid dynamics are more strongly dominated by niche and local geography, rather than phylogeny or season. Our results highlight the diversity of the AMR reservoir in these species and niches, and the importance of local strategies for controlling the emergence and spread of AMR.

---

*Enterobacteriaceae* can persist across diverse environmental niches^10^ and also cause clinical infections, with AMR in *Enterobacteriaceae* emerging as a major problem in the last decade^11,12^. Dissemination of AMR genes often occurs via mobile genetic elements (MGEs) which can transfer intra- and inter-species, both locally^13^ and globally^14^. Freshwater, wastewater and livestock-associated strains of *Enterobacteriaceae* have been proposed as reservoirs for AMR genes in clinical isolates^15–18^, but the links between these remain cryptic^19^. Current understanding of the ecology and evolution of pangenomes is incomplete^20^, with ongoing debate about the roles of niche adaptation and selection^21–24^. Published *Enterobacteriaceae* genomes are biased towards clinical isolates, sampling frames reflecting truly interlinked communities are limited, and much remains unknown about the population genetics of *Enterobacteriaceae*^25^ and the role of plasmids in non-clinical contexts^26^.

Genomic studies of *Enterobacteriaceae* have predominantly used short-read whole genome sequencing (WGS). AMR genes and their flanking regions are frequently fragmented in shortread assemblies due to repetitive elements and structural rearrangements^13,27^. Combining short- and long-reads (‘hybrid assembly’) produces complete, high-quality genomes^28^, allowing accurate structural resolution. Here, we used hybrid assembly of 828 sympatric *Enterobacteriaceae* (*Citrobacter*, *Enterobacter, Escherichia* spp., and *Klebsiella*) to characterise the pangenome of these genera considering both niche (cattle, pig, sheep, or wastewater treatment works (WwTW)-associated) and geography (sampling location).

## A diverse collection of complete genomes from livestock and water-borne niches

We collected samples from nineteen locations ≥5 km apart (maximum distance: 60km) in South-central England (United Kingdom) in 2017, namely: fourteen livestock farms (four pig, five cattle, five sheep) and water sources around five WwTWs over three seasonal timepoints (Fig. 1a). A subset of 832/2098 cultured isolates from pooled samples from each sampling location underwent short- and long-read sequencing and hybrid genome assembly (Fig. 1b, see Methods), resulting in 828 high-quality genomes (Table S1: *n*=496 from livestock farms, *n*=332 from WwTWs), from four genera: *Citrobacter* (*n*=128), *Enterobacter* (n=71), *Escherichia* (*n*=553), and *Klebsiella* (*n*=76). Most farm isolates were *Escherichia* spp. (451/496, 90.9%), with WwTW isolates having roughly even proportions of genera (Fig. S1). Isolates contained a median of 1 AMR gene (range: 0-23); *Klebsiella* isolates carried a median of 4 (range: 1-18).

**Figure 1.**
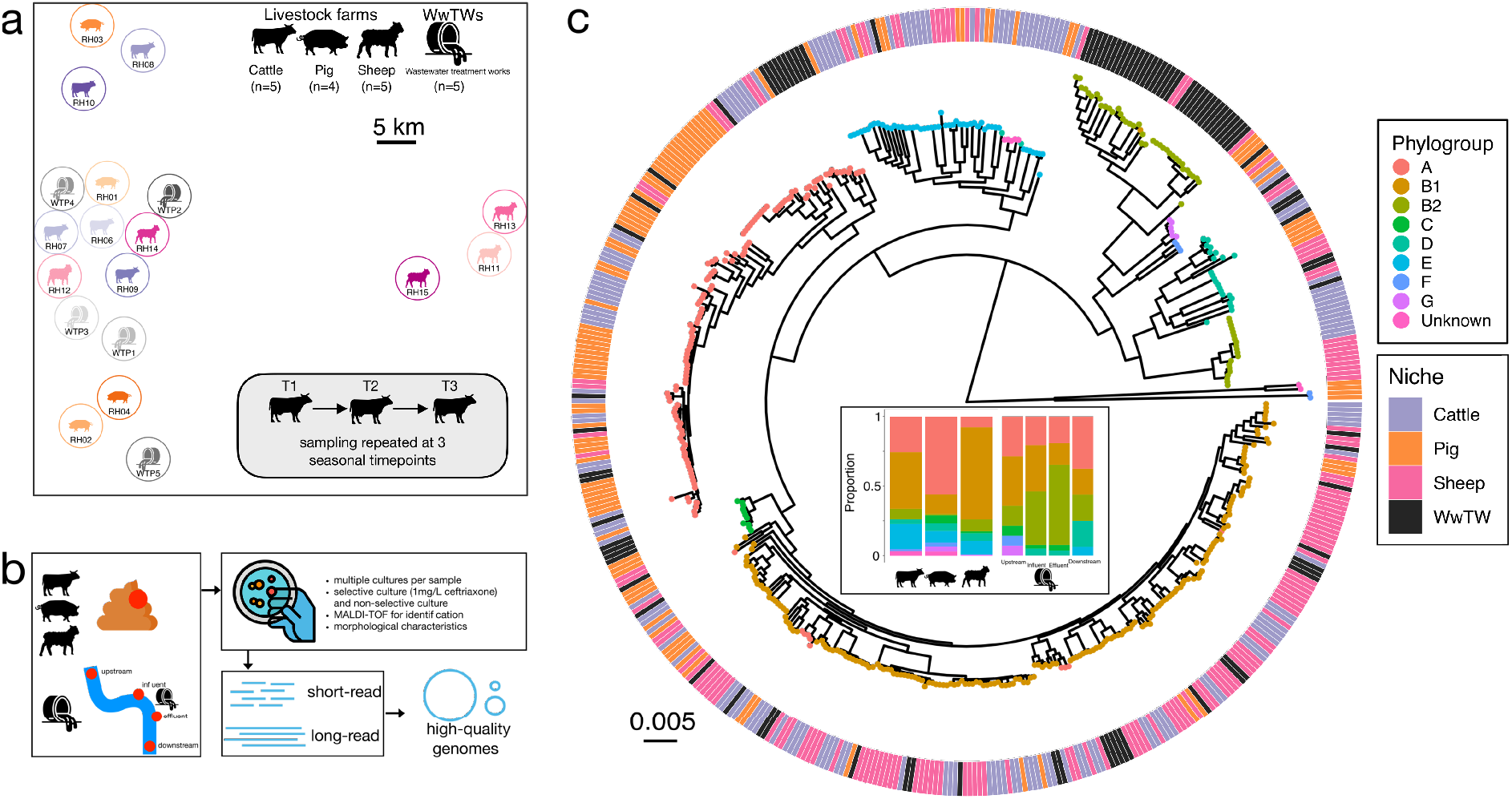
Overview of the diverse *Escherichia coli* isolates in this study. **(a)** Relative sampling locations of the farms (cattle, pig, sheep) and wastewater treamtent plants (WwTWs) in this study, sampled at three different timepoints. **(b)** Schematic illustration of the sampling, culture and sequencing workflow, resulting in high-quality genome assemblies with a median of 1 circularised chromosome and 2 circularised plasmids per assembly. **(c)** Mid-point rooted core genome phylogeny of *E. coli* isolates (*n*=488) based on, with tips coloured by phylogroup and ring colours showing sampling niche. Inset panel at centre of phylogeny shows phylogroup abundances (as proportion of isolates) from different sampling niches.

Isolates were highly diverse, containing novel diversity not present in published genomes (Fig. S2). *Escherichia* diversity included all main *E. coli* phylogroups, 53 *E. fergusonii*, and 13 isolates from clades I, II, III, and V (Fig. 1c). Phylogroup B2 was strongly associated with WwTWs compared to livestock (34.3% vs. 5.1% of *Escherichia* isolates) particularly in influent and effluent samples (Fig. 1c). Pigs had a greater proportion of phylogroup A isolates (Fig. 1c). Of 187 identified *E. coli* multilocus sequence types (STs), 56.1% (105/187) were seen only once, similar to the 61% observed by Touchon et al. in a study of non-clinical *E. coli*^29^. Only 12 *Escherichia* STs were seen in both livestock and WwTW isolates, with phylogroup B1 the most represented (5/12 STs). ST10 was the most prevalent ST (*n*=45), seen in 10/14 farms and 3/5 WwTWs.

Considering only livestock *E. coli* isolates, over time, there was a persistent phylogroup signature of both livestock host and farm, with individual farm explaining more variance than livestock type (*R^2^* 28.1% vs. 25.5%, Fig. S3). However, livestock type explained less variance for STs than phylogroups (*R^2^* 8.5%), with only 39/131 STs (29.8%) seen on more than one farm. There were only 26 instances where an *E. coli* ST was observed over time on the same farm (involving 16 STs) and the majority of these (22/26) were STs also seen across farms (Fig. S4). Considering *E. coli* strain clusters using a core genome distance of <100 single-nucleotide variants (SNVs) (maximal diversity observed across sampled *E. coli:* 211,251 SNVs; median pairwise distance 46,144 SNVs), there were 280 isolate pairs with <100 SNVs, of which 181 (64.6%) were isolates cultured from the same pooled sample (i.e. same farm, same timepoint) (Fig. S5a). Overall, 10.5% of all isolate pairs from the same pooled sample had <100 SNVs between them, compared to 1.4% (*n*=52) of isolate pairs from different timepoints on the same farm and 0.2% (*n*=44) between different farms of the same animal (Fig. S5b). Notably, of the latter, 41/44 were between cattle farms, and 36 involved a single cattle farm (RH06). There were only three isolate pairs with <100 SNVs between farms of different animals (Fig. S5a).

Notably, all of these were between farms in close geographic proximity (two instances from pig farm RH03 and cattle farm RH10, one instance from cattle farm RH07 and sheep farm RH12; see Fig. 1a for distances), suggesting local strain movement. Taken together, this indicates that different livestock hosts have a stable balance of *E. coli* phylogroups but each farm harbours substantial strain-level diversity. There were no isolate pairs with <100 SNVs between WwTW and livestock niches, and only three isolate pairs occurred across timepoints at WwTWs (all at a single WwTW).

## Plasmid gene repertoires are linked to genus and niche

We recovered 2,293 circularised plasmids across all *Enterobacteriaceae*, ranging in size from 1,240-824 kbp (median: 43 kbp; Table S2). There were 298/2,293 (13.0%) with no identifiable plasmid replicon and the majority of these were from WwTW isolates (192/298, 64.4%). Multiple replicons were carried by 723/2,293 (31.5%) and these plasmids tended to be larger (median length: 106,811 bp vs. 6,275 bp for single replicon plasmids). Of *E. coli* isolates with complete genomes, over two thirds (70.4%, 245/348) carried a plasmid with an IncFII replicon. 43.0% of circularised plasmids (986/2,293) had at least one match with >99% identity to other publicly available plasmid sequences (Fig. S2b). However, 12.3% (282 of 2,293) had a top identity score of <95% to a previous known sequence (Fig. S2b), and 17 plasmids with no match were identified, suggesting novel plasmid diversity in our setting. We grouped circularised plasmids into 611 distinct plasmid clusters, which closely matched gene content (Fig. S6a). The synteny of shared genes was strongly conserved, supporting the concept of plasmid ‘backbones’ (Fig. S6b).

A median of 3.3% of genes were on plasmids (range: 0-16.5%), with substantial variation by genus and niche (Fig. S7a). Accounting for plasmid copy number, *E. coli* isolates had a median of 5.7% of DNA present on plasmids, which was substantially higher in pig farm isolates (median: 10.1%; Fig. S7b). Chromosomal genes were highly genus-specific (*R^2^*=55.0%); the plasmid-borne pangenome was far more variable but still genus-specific (*R^2^*=6.5%) (Fig. 2). Within *E. coli*, plasmid gene content was linked to niche (*R^2^* 5.6%) and phylogroup (*R^2^* = 5.2%), with a stronger interaction between niche and phylogroup (*R^2^*=7.9%) (Fig. 2). Non-mobilizable plasmid clusters were less commonly shared between different phylogroups within farms compared to mobilizable or conjugative plasmids (Fig. S8). Although AMR genes were predominantly found in conjugative/mobilizable plasmid clusters, plasmid clusters with AMR genes were not more commonly distributed across multiple phylogroups (Chi-squared test *χ^2^*=0.64, *p* = 0.42; Fig. S8). On pig farms however, the majority of conjugative plasmid clusters across multiple phylogroups carried AMR genes, suggesting an important role within this niche.

**Figure 2.**
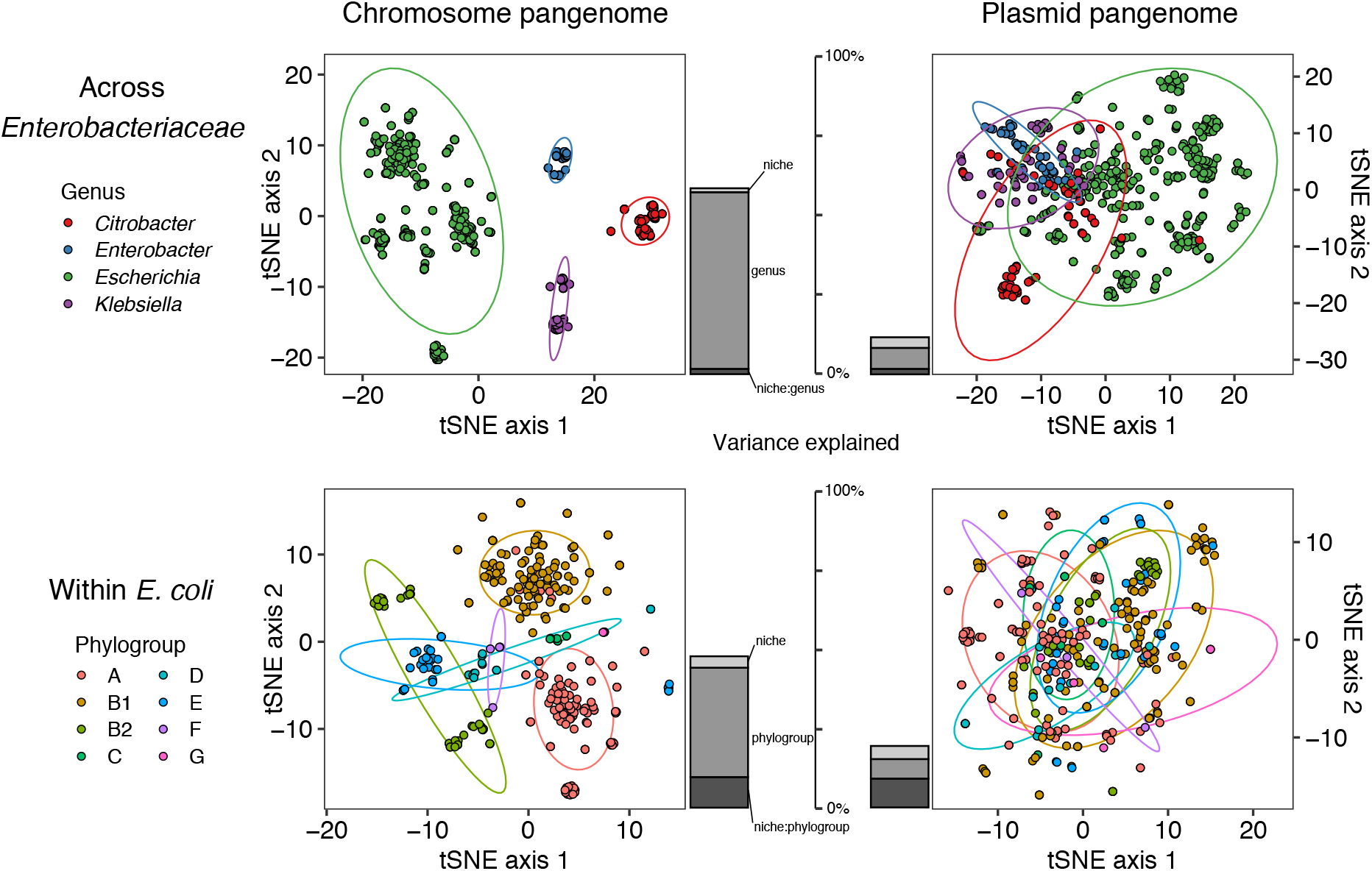
The plasmid-borne component of the pangenome is structured by niche and phylogeny, with greater variation than in the chromosomal component. Plots are shown for all isolates in four genera across *Enterobacteriaceae* (top row) and for *E. coli* (bottom row), for both the chromosomal component of the pangenome (left column) and the plasmid-borne component (right column). Stacked bar charts show the variance in gene content explained by niche, phylogeny (genus or phylogroup) and their interaction. The plasmid-borne component has greater residual variance than the chromosomal component, with a comparatively stronger niche-phylogeny interaction (darkest shaded bar).

Positive epistasis between large (>10 kbp) and small plasmids has been suggested to promote plasmid stability in *Enterobacteriaceae*^30^. In *E. coli* isolates with complete genomes (*n*=348), we observed a significant association between small and large plasmid presence (Chi-squared test *χ*^2^=4.44, *p*=0.035), with 45.7% carrying at least one large (>10 kbp) and one small plasmid and only 3.7% carrying a small plasmid without a large plasmid. We also found evidence of specific plasmid-plasmid associations. For example, cattle *E. coli* isolates showed co-occurrence of a ColRNA plasmid (cluster 37: median length 4.6 kbp) and an IncFII plasmid cluster (cluster 279: median length 106 kbp), with 14/16 isolates with the ColRNA plasmid also carrying the larger IncFII plasmid. Isolates were from three phylogroups (A: *n*=2, B1: *n*=5, E: *n*=9) and four farms, suggesting a robust association which reflects plasmid epistasis independent of chromosomal background.

## Plasmids carry an over-representation of AMR genes and insertion sequences

Plasmids carried more diverse and less genus-restricted genes. Despite carrying just 3.3% of total gene content, plasmid-borne genes accounted for 11.5% of unique genes (8.9-17.0% considering each genus; Fig. S9) and 40.1% were seen in more than one genus (19.6-55.6% considering each genus; Table S3). Plasmids also had a much greater burden of AMR genes: considering isolates with circularised chromosomes (see Methods), 901/1,876 AMR genes (48.0%) were found on plasmids i.e. a 14.5x relative burden in plasmids. Of 26,565 insertion sequences (ISs), 3,695 (21.7%) were found on plasmids (6.6x relative burden). There was a weak correlation between the number of plasmid- and chromosome-associated AMR genes within an isolate (Spearman’s *ρ*=0.11, *p*=0.004) but a strong positive correlation for the number of ISs (Spearman’s *ρ*=0.40, *p*<0.001) (Fig. S10a), seen across genera (Fig. S10b).

We observed different patterns of ISs across chromosomes and plasmids (Fig. S11). Some ISs were strongly associated with plasmids, the strongest association being for *IS26*. However, 27.5% of isolates carrying IS*26* on a plasmid also carried it on their chromosome, consistent with its extremely active behaviour^31^. The most prevalent IS on both chromosomes and plasmids was IS*Kpn26*, with 50.2% of IS*Kpn26*-positive isolates having it both chromosomally and plasmid-borne. Considering *Escherichia*, WwTW isolates showed a greater diversity of ISs, with 65% of ISs found in a higher proportion of WwTW isolates compared to those from farms (Fig. S12), including IS*30* which has been proposed as a marker for naturalized wastewater populations of *E. coli*^32^. Overall, ISs had random levels of co-occurrence on *Escherichia* plasmids (upper-tail *p*=0.85 from null model simulations of checkerboard score, see Methods; Fig. S13a), suggesting that ISs frequently move independently between plasmid backgrounds. Contrastingly, AMR genes significantly co-occurred (upper-tail *p*=0.02; Fig. S13b), suggesting co-selection on plasmids.

## Plasmids carrying AMR genes show features suggestive of selection

Plasmids fell into two broad classes across genera: small multicopy plasmids (<10 kbp, 10-100X copy number) and large low-copy plasmids (>10 kbp, <10X) (Fig. 3a). AMR plasmids were almost all large low-copy plasmids (173/184, 94.0%). Overall, plasmids had a lower relative GC-content than their host chromosomes (median difference −2.5%, Fig. 3b), and plasmids predicted to be mobile had a smaller relative difference. However, this difference was less marked for AMR plasmids (median −0.3%) across mobility categories (Fig. 3b). Nearly half had a higher GC-content than their host chromosome (46.7% vs. 17.7% of non-AMR plasmids), suggesting AMR plasmids are under selective pressure to maintain their function.

**Figure 3.**
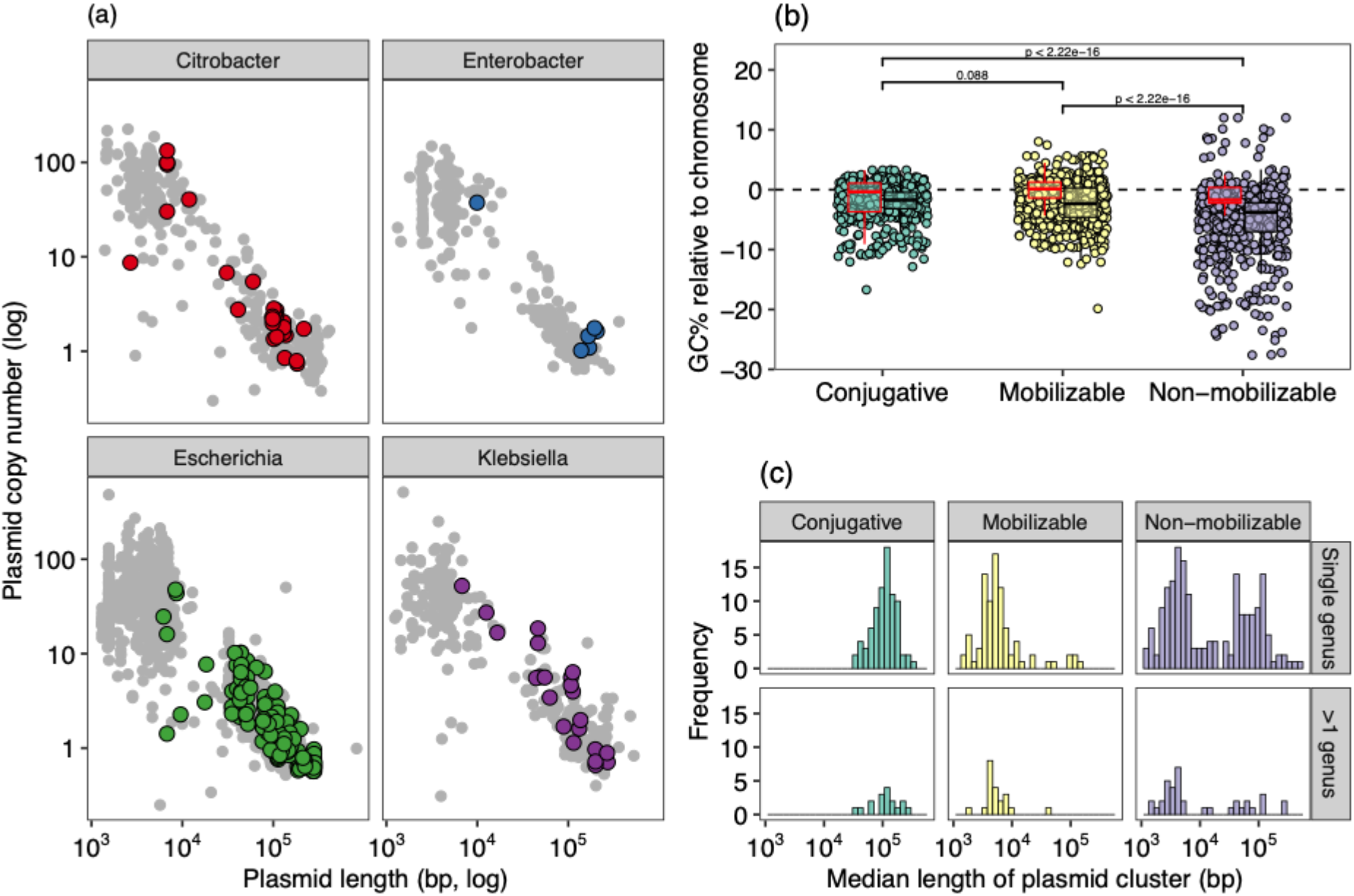
Distinct plasmid lifestyles between AMR and non-AMR plasmids. **(a)** Plasmid length (x-axis) and inferred copy number (y-axis) of all circularised plasmids (*n*=2,293), faceted by genus. Plasmids with ≥1 AMR gene (coloured points) tended to be larger and present in lower copy numbers. **(b)** Relative GC-content of all plasmids to their host chromosome for all circularised plasmids present in an assembly with a circularised chromosome (*n*=1,753 plasmids across 616 isolates), split by predicted plasmid mobility. Boxplots are shown for plasmids with ≥1 AMR gene (red) or no AMR genes (black). Comparisons with *p*-values are shown for all plasmids within a predicted mobility class. **(c)** Length distributions of plasmid clusters (see Methods).

## Evidence for recent horizontal gene transfer across genera and within isolates

We identified 2,364 potential horizontal gene transfer (HGT) events involving transfers of sequence >5,000 bp between isolates of different genera (see Methods). Isolates from the same farm were ~10x more likely to show evidence of cross-genera HGT than would be expected (Chi-squared test *χ^2^*=1159, *p*<0.001; Fig. S14), and 12.3% of these cross-genera HGT events involved at least one AMR gene, with most of these AMR HGT events between pig isolates (37/48, 77.0%). Movement of genes can also occur within genomes. We therefore also investigated occurrences where the same gene was present on both the chromosome and plasmid(s) within an *E. coli* genome. We observed distinct differences between niches, with increased amounts of chromosome-plasmid sharing in pig and WwTW isolates compared to cattle and sheep (Fig. S15).

## Quantifying the roles of phylogeny, niche and geography in the *E. coli* pangenome

To understand the strength of different factors shaping the pangenome, we analysed the pangenome of *E. coli* in more detail. Isolates recovered from the same location spanned total *E. coli* diversity (Fig. 4a). Inter-isolate core genome distances were strongly correlated with chromosomal gene repertoire relatedness (GRR) (Fig. 4a). Core genome distance explained the majority of variance in chromosomal GRR (Fig. 4b), but there was a consistent contribution from geography and time: isolates from the same pooled sample sharing more genes than would be expected (+1.2%), as did isolates from the same farm at different timepoints (+0.5%) (Fig. 4b). There was no such effect for isolates from different farms of the same livestock, suggesting this reflects local geography rather than adaptation to livestock host. Although the variance explained was much lower, local geography effects were also observed for plasmid GRR (Fig. 4c), but core genome distance was uncorrelated with plasmid GRR apart from for near-identical strains (Fig. 4d). Isolates from different STs from different farms of the same livstock could still have high plasmid GRR (Fig. 4e), suggesting that host-specific plasmids may facilitate niche adaptation.

**Figure 4.**
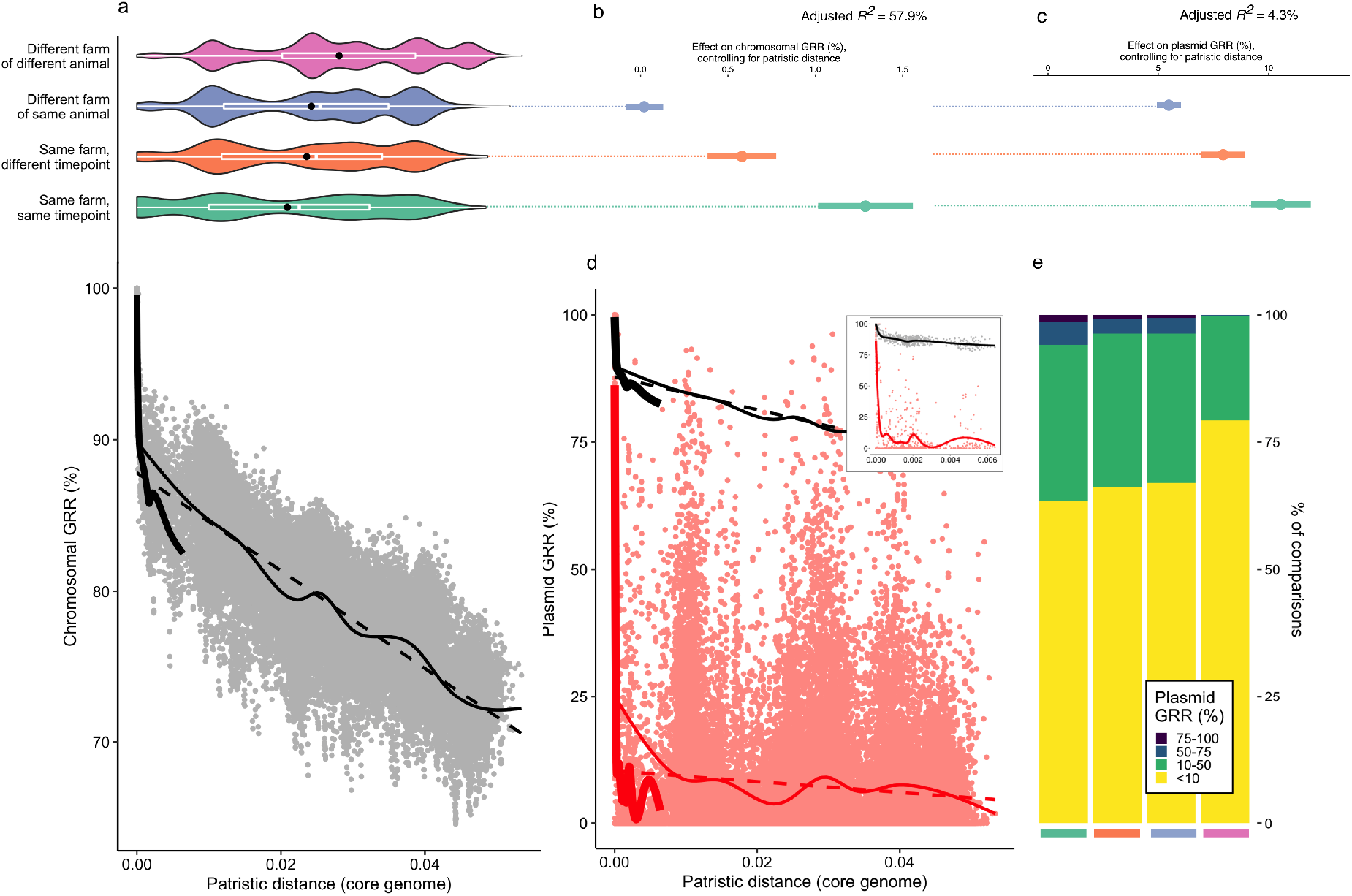
The interplay of phylogeny and niche in the *E. coli* pangenome. **(a)** Pairwise comparisons of gene repertoire relatedness (GRR) for chromosomal genes show that chromosomal GRR falls off rapidly at small patristic distances, followed by an approximately linear decrease. Fits show intra-ST comparisons (thick black line), all comparisons (thin black line), and a linear model (dashed black line). Violin plots above show the distribution of patristic distances depending on the relative sample source of the two isolates in the pairwise comparison (white boxplot: median and IQR; black point: mean), showing that even isolates cultured from the same sample (same farm, same timepoint) span equivalent diversity to isolates cultured from different locations. **(b)** Coefficients from a linear model for chromosomal GRR with an interaction term with patristic distance (excluding intra-ST comparisons). **(c)** Variance explained by phylogeny and geography for chromosomal and plasmid GRR. **(d)** GRR for plasmid-borne genes with patristic distance. Fits show intra-ST comparisons (thick red line), all comparisons (thin red line), and a linear model (dashed red line). Inset panel shows left-hand region of the plot with only intra-ST comparisons, with chromosomal GRR relationship also shown (grey points, black line). **(e)** Plasmid GRR comparisons shown by isolate sources, excluding intra-ST comparisons. Colours on x-axis are the same as in (a). Plots include all *E. coli* isolates with a circularised chromosome (*n*=363).

## Conclusions

We have investigated the pangenome of major genera of sympatric *Enterobacteriaceae* from locations within a 30km radius, using a diverse set of non-clinical isolates cultured from the same samples, and focusing in detail on *E. coli*. Despite high overall diversity, with the majority of strains only observed once in the dataset, we observed the persistence of strains and plasmids on farms over the course of a year. Our results highlight the combination of persistence and dynamism that characterises *Enterobacteriaceae* genomes at multiple scales, with relevance both for understanding the population structure of species within *Enterobacteriaceae* and for managing AMR. The existence of farm-level differences in *E. coli* populations which persist over time, with a small number of possible inter-farm transfers, suggests that livestock farms function as distinct but linked niches. It could be that “everything is everywhere” (frequent movement of strains and genes between farms) but “the environment selects” (different farms have different selective pressures). However, the observation of persistent strains over the course of a year on farms, despite presumably varying selective conditions, and the overrepresentation of putative cross-genera HGT events in isolates at the same location suggests that geographical effects or intrinsic properties of certain bacterial/MGE lineages could affect the evolution of AMR on such timescales. Future modelling work and investigation will be required to distinguish these hypotheses. Overall, our findings underline the importance of local control strategies for the emergence and spread of AMR beyond clinical settings.

Resource limitations meant that we were unable to sequence and genetically evaluate all isolates that were cultured, and despite our detailed sampling we will not have captured all the persistence, HGT and strain sharing events across niches. Although this study is unprecented in evaluating four genera in such detail, AMR gene dissemination and important structural associations of AMR genes and MGEs may also be occurring within other genera not studied here. Furthermore, we did not investigate the relationship between isolates in this study and clinical human compartments in the same study area; this is ongoing work.

In conclusion, our study highlights the plastic and dynamic nature of AMR gene dissemination within the pangenome of major *Enterobacteriaceae* in several important non-clinical niches. It also demonstrates how robustly evaluating the flow of AMR genes and MGEs across highly diverse and dynamic niches is challenging even with extensive sampling. The implications of this for adequately understanding dissemination and selection of AMR genes in a ‘One Health’ context should not be under-estimated.

## Methods

Isolates were sequenced from samples collected as part of the REHAB project in 2017, which aimed to characterise non-clinical *Enterobacteriaceae* populations in four different niches within a defined study area of South-central England: cattle farms, pig farms, sheep farms, and water environments linked to wastewater treatment works (WwTWs). Sampling occurred at each location at three separate timepoints (TPs): January-April 2017 (TP1), June-July 2017 (TP2), October-November 2017 (TP3).

### Farms

Five cattle farms, five sheep farms and four pig farms were recruited from the study area following a defined recruitment process (described in more detail in AbuOun et al.^33^). Briefly, we aimed to recruit the five largest farms for each livestock type within the area using local APHA databases, progressively inviting the next largest farm if a farm declined. All participating farmers provided written consent for farm sampling for research purposes and farm samples were taken between January and November 2017 on three separate visits (‘timepoints’) for each farm. Each farm was divided in five or fewer ‘epidemiological groups’, defined as a group of animals expected to share similar characteristics and managed in the same way. Ten pooled samples were collected from each of these groups, with each sample composed of small pinches of fresh faeces from the floor combined into a small composite sample around 5cm in diameter. Each group’s ten samples were pooled, diluted up to 10^-5^ in phosphate buffer solution (PBS) (pH 7.2) and plated on to CHROMagar™ ECC (CHROMagar Microbiology, Paris, France) and CHROMagar™ ECC plates containing 1mg/L cefotaxime as a marker for multi-drug resistance. Up to ten colonies were collected from 1mg/L cefotaxime-supplemented plates and fourteen colonies from CHROMagar™ ECC plates; where ten colonies were not recovered, additional colonies were taken from the CHROMagar™ ECC plates, resulting in 24 isolates per farm. Pure isolate sub-cultures were subsequently stored at −80°C in MicroBank beads (Pro-Lab Diagnostics, Neston, Cheshire, UK), and the bacterial species confirmed using MALDI-TOF (Bruker, Coventry, UK) or 16S rRNA sequencing^34^. The median number of sequenced isolates for a farm-timepoint combination was twelve (range: 9-14), with 496 farm isolates in total: cattle (*n=*178), pig (*n*=144), sheep (*n*=174).

### Wastewater treatment works (WwTWs)

Five WwTWs were selected based on a defined recruitment process (described in more detail in Read et al.^35^) including; geographic location within the study area, wastewater treatment configuration, wastewater population equivalent served, consented flow, and the accessibility of the effluent receiving river for sampling both upstream and downstream. Sampling took place in 2017 over three sampling rounds: February–March (TP1), June–July (TP2), and October–November (TP3). Sewage influent samples were collected after WwTW coarse screens and effluent samples were collected at the last sampling point before entering the river. For each sampling round, ~6 repeated 200 ml samples of influent and effluent were collected between 9 am and 12 pm, using a sampling pole and sterile Whirl-Pak collection bags. Repeat samples in each round were pooled prior to processing, to reduce the impact of temporal variability in wastewater flows and composition. Sediment samples were collected from 100 m upstream and 250 m downstream of the effluent entry point into the river. Sediment samples were collected using a custom sampling pole that held a removable 50 ml plastic centrifuge tube (Sigma, UK). Using a fresh sterile 50 ml tube each time, sediment from the riverbed was collected from the surface layer at three points at each sampling site; near bank, the centre of the river, and the far bank. These samples were pooled prior to analysis to account for spatial variability in sediment composition. Influent, effluent and sediment samples were stored in an insulated box at ~4 °C until getting back to the laboratory (<6 h). Influent, effluent, 100 m upstream and 250 m downstream environmental samples collected from each sewage treatment works were transferred to the laboratory on ice and processed within 24 hours of collection. Each sample was vortexed briefly, serial diluted to 10^-3^ in nutrient broth containing 10% glycerol (Oxoid, Basingstoke, UK) and plated on to CHROMagar™ Orientation agar (Chromagar, Paris, France) and CHROMagar™ Orientation agar supplemented with 1 μg/ml cefotaxime (Cambridge Biosciences, Cambridge, UK). Colonies with putative morphology for species of interest were subcultured from dilution plates with suitably isolated growth. A total of up to 20 colonies was picked per sample: up to ten colonies were picked from the 1mg/L cefotaxime-supplemented plates and the remainder picked from the non-supplemented plates. Pure isolates subcultured on Columbia blood agar (CBA) (Oxoid, Basingstoke, UK) were subsequently stored at −80°C in nutrient broth containing 10% glycerol, and bacterial species confirmed using MALDI-TOF (Bruker, Coventry, UK).

### DNA sequencing

A subset of isolates were selected for sequencing to represent diversity within the four major genera within each niche, including the use of third-generation cephalosporin resistance as a selective marker to identify a sub-group of multi-drug resistant isolates within each genus. 832 isolates were each sequenced with both a short-read (Illumina HiSeq 4000) and a long-read sequencing approach (four isolates selected for sequencing failed subsequent hybrid assembly and were not included in further analyses). For the first timepoint, the latter involved sequencing using either PacBio SMRT (*n*=192) or Oxford Nanopore Technologies (ONT) methodologies (*n*=127). The results of a pilot study comparing sequencing and assembly approaches using a subset of REHAB isolates^28^ were used to inform the choice of ONT as the long-read sequencing approach for all isolates from the second (*n*=255) and third (*n*=254) timepoints.

Isolate stocks from −80°C storage were cultured on to CBA and supplemented with cefpodoxime (Fisher Scientific, USA) 10 μg discs for isolates not sensitive to cefotaxime during original sample isolation. DNA was extracted using the Qiagen Genomic tip/100G (Qiagen, Venlo, Netherlands) according to the manufacturer’s instructions. DNA concentration was quantified by Qubit^®^ 2.0 fluorimeter (Invitrogen, UK), and quality and fragment size distribution assessed by TapeStation 2200 (Agilent, Santa Clara, USA). ONT sequencing libraries were prepared by multiplexing 6-8 DNA extracts per flow cell using kits SQK-RBK004, SQK-LSK108 and EXP-NBD103 according to the manufacturer’s protocol. Libraries were loaded onto flow cell versions FLO-MIN106 R9.4(.1) SpotON and sequenced for 48 h on a GridION (ONT, Oxford, UK).

### Genome assembly

We used the hybrid assembly and sequencing methods described in our pilot study^28^ to produce high-quality *Enterobacteriaceae* genomes from short and long reads. In brief, we used Unicycler (v0.4.7)^36^ with ‘normal’ mode, --min_component_size 500, --min_dead_end_size 500, and otherwise default parameters. Final assemblies had a median of four contigs (IQR: 3-8, range: 1-391), with a median of two circularised plasmids (IQR: 1-4, range: 0-14). The majority (616/828, 74.4%) of assemblies had a circularised chromosome, and 558/828 (67.3%) were complete i.e. chromosome and all plasmids circularised (Table S1).

### Genome assignment and typing

We assigned species and sequence type (ST) from assembled genomes using mlst v2.16.4^37^. We also validated species assignments by downloading all NCBI Refseq complete genomes for the four genera under study as of June 4 2020 and using fastANI (v1.3)^38^ to compute average nucleotide identity scores against reference genomes for each assembled genome. We took the species assignment of the top hit for each assembled genome. Furthermore, we manually checked genus assignments using a tSNE plot of isolate genomes against a collection of reference genomes (not shown) and made corrections to the assignment if necessary. We used ClermonTyping (v1.4.1)^39^ to assign phylogroup to *n*=553 *Escherichia* isolates. Considering the genus *Escherichia*, there were 553 isolates, 410 with circularised chromosomes, and of these 379 were complete genomes containing 961 complete plasmids in total. Considering only *E. coli*, there were 502 *E. coli* isolates, 372 with circularised chromosomes, and of these 348 were complete genomes containing 878 complete plasmids in total. A minority of genomes were *E. fergusonii* (n=51), from clades I-V (*n*=14), or could not be typed (*n*=7), with *n*=481 genomes from within the principal *E. coli* phylogroups (A: *n*=131, B1: *n*=193, B2: *n*=59, C: *n*=11, D: *n*=25, E: *n*=50, F: *n*=6, G: *n*=6).

Sequenced isolates from three other *Enterobacteriaceae* genera included: *Citrobacter* (*n*=128: 82 *C. freundii* and 46 unassigned *Citrobacter sp*.), *Enterobacter* (*n*=71: 59 *E. cloacae* and 12 unassigned *Enterobacter sp*.); and *Klebsiella* (*n*=76: 40 *K. pneumoniae*, 30 *K. oxytoca*, 2 *K. aerogenes*, and 4 unassigned *Klebsiella sp*.). The majority of farm-associated isolates were *E. coli*, whereas WwTW-associated isolates had roughly equal numbers of genera (Fig. S1). This reflects both the diversity present in each niche and the selection strategy to sequence equal numbers across genera where feasible.

### Pangenome analysis

All genomes were annotated with Prokka (v1.14.0)^40^. Genes were clustered into gene groups using Roary (v3.12.0)^41^ across all isolates at various sequence identity thresholds with the maximum number of clusters set to 300,000 (-g 300000) and without splitting paralogs (-s). At a 95% identity for blastp, there were 139,788 gene groups across all genera. Further to this analysis, genes were also clustered at a higher sequence identity (>99% identity threshold) in order to identify recent HGT events, which gave 214,743 gene groups across all genera. For *n*=616 isolates with circularised chromosomes, we split the genome into chromosomal and plasmid-borne components (i.e. all other contigs) to analyse the genomic location of genes. We excluded isolates without circularised chromosomes from this analysis. For *n*=488 E. *coli* isolates (excluding *E. fergusonii* and clades I-V), we used Panaroo (v0.1.0)^42^ to extract a core genome alignment based on 2,915 concatenated core genes (Fig. 1c). The phylogeny was produced using iqtree (v1.6.11)^43^, with branch lengths not corrected for recombination, and plotted with ggtree (v2.0.1).

### Plasmid annotation and clustering

We searched all plasmids against PLSDB (version: 2020-03-04)^44^ which contains 20,668 complete published plasmids, using ‘screen’ in mash (v2.0)^45^ and keeping the top hit. All plasmids had a match apart from 17 small plasmids predicted to be non-mobilizable (median length 4.8 kbp, range 2.9-20.7 kbp), from *Escherichia* (n=11), *Enterobacter* (n=2) and *Citrobacter* (n=4). We clustered plasmids using mob cluster and assigned replicon types with mob typer, both part of the MOB suite^46^. Mob cluster uses single linkage clustering with a cutoff of a mash distance of 0.05 (corresponding to 95% ANI), resulting in 611 clusters (Table S2). In total, there were 134 different combinations of replicons observed on plasmids (‘replicon haplotypes’). The most abundant replicon was IncFIB (n=460) which was seen across all niches (pig [n=81], cattle [n=113], sheep [n=78], and WwTWs [n=188]). Only nine small multicopy plasmids (~6 kbp) carried AMR genes, all of which had a ColRNAI replicon; such ColRNAI plasmids have been proposed to be sources of evolutionary innovation^47,48^.

We considered the relationship between such ‘distance-free’ clustering and plasmid gene content. Based on gene clustering with Roary (see above), we compared the structure of circularised plasmids using all connecting edges between two genes. We defined the resemblance for both gene content (gene presence/absence) and gene structure. The gene content resemblance between two plasmids with *n*_1_ and *n*_2_ genes respectively, with *N* genes in common, was defined as *r*_content_=2*N*/(*n*_1_+*n*_2_). The edge structure resemblance between two plasmids with *g* gene-gene edges in common, was defined as *r*_edge_=2*g*/(*n*_1_+*n*_2_). Typically *r*_edge_<*r*_content_ but this definition does allow for the case where repeated genetic elements produce *r*_edge_>*r*_content_ (e.g. Fig. S6b).

### Comparison of plasmid-borne and chromosomal pangenome components

To visualize cross-genera pangenomes (e.g. Fig. 2), we used t-distributed Stochastic Neighbor Embedding (t-SNE). We used the Rtsne function with a perplexity of 30 on gene presence/absence matrices using the Rtsne R package. To conduct permutational analyses of variance, we used the adonis function from the vegan R package on the matrix of pairwise Jaccard distances, which was calculated using the vegdist function. For between-genera analyses, we used the formula *dist~niche*genus*. For within-*Escherichia* analyses, we used the formula *dist~niche*phylogroup*.

### Detection of antimicrobial resistance genes and insertion sequences

We searched assemblies using ABRicate (v0.9.8)^49^ for acquired resistance genes (i.e. excluding mutational resistance) in the the NCBI AMRFinder Plus database (PRJNA313047). We used a minimum identity threshold of 90% and a minimum coverage threshold of 90% (Table S4). Isolates cultured selectively from cefotaxime-supplemented plates carried more AMR genes than non-selectively cultured isolates (median of 7.5 vs. 1.0), as expected. We also searched for insertion sequences (ISs) using the ISFinder database^50^ as a database in ABRicate with the same identity and coverage thresholds (Table S5).

### Detection of recent horizontal gene transfer events

We performed an all-against-all comparison of assemblies with mummer (v3.23-2)^51^ using the-maxmatch option to identify shared sequences of length >5,000 bp between genomes of different genera (these could include both transfer of whole plasmids or partial sequences). For comparing the observed distribution of cross-genera HGT events to the expected, we assumed a random distribution drawn from all possible cross-genera comparisons from livestock isolates.

### Distribution of insertion sequences

We constructed the bipartite presence/absence network of ISs and replicon haplotypes for the 34 replicon haplotypes which were observed on 10 or more plasmids. We simulated null models of co-occurrence patterns using the cooc_null_model with null model sim9, which fixes the row and column sums of the presence/absence matrix, in the R package EcoSimR (v0.1.0)^52^. Simulations used n=10,000 iterations with a burn-in of 500 iterations.

### Modelling of gene repertoire relatedness (GRR)

We selected a subset of *E. coli* genomes with a circularised chromosome (*n*=363) and used the core genome tree constructed with iqtree (Fig. 1c, dropping other *E. coli* isolates) to calculate patristic distances between isolates. We calculated chromosomal and plasmid GRR for all pairwise comparisons using output from roary (95% identity threshold, as above) and fit linear models for GRR (Fig. 4).

## Data availability

Sequencing data and assemblies have been uploaded to NCBI under BioProject accession PRJNA605147. Biosample accessions for all isolates are provided in Table S1.

## Image credits

The following images are used in Figure 1: Petri dish icon made by monkik; pig, cow, sheep and faeces icons made by Freepik; WwTW symbol made by Smashicons (all sourced from flaticon.com).

## Acknowledgements

The REHAB consortium is represented by the following: Manal Abuoun, Muna F. Anjum, Mark J. Bailey, Howard Brett, Mike J. Bowes, Kevin K. Chau, Derrick W. Crook, Nicola de Maio, Daniel Gilson, Sophie George, H. Soon Gweon, Alasdair T. M. Hubbard, Sarah J. Hoosdally, James Kavanagh, Hannah Jones, William Matlock, Tim E. A. Peto, Daniel S. Read, Robert Sebra, Liam P. Shaw, Anna E. Sheppard, Richard P. Smith, Emma Stubberfield, Nicole Stoesser, Jeremy Swann, A. Sarah Walker, Neil Woodford.

## Author contributions

Using the CRediT system, author contributions were as follows: conceptualization (LPS, MJB, DWC, DSR, MFA, ASW, NS), methodology (LPS, ASW, NS), software (LPS, JS), validation (LPS, KKC, JK, MA, ES, LB, GR, ATMH, HP, RS, NS), formal analysis (LPS), investigation (KKC, JK, MA, ES, HSG, LB, GR, MJBo, ATMH, HP, JS, DG, RPS, RS, NS), resources (all authors), data curation (LPS, KKC, JK, MA, ES, LB, GR, ATMH, HP, JS, DG, RPS, RS, NS), writing - original draft (LPS, ASW, NS), writing - review & editing (all authors), visualization LPS), supervision (TEAP, MJBa, DWC, DSR, MFA, ASW, NS), project administration (MA, RPS, SJH, DSR, MFA, ASW, NS), funding acquisition (MJB, DWC, DSR, MFA, NS).

## Competing interest declarations

The authors declare no competing interests.

## Funding

This work was funded by the Antimicrobial Resistance Cross-council Initiative supported by the seven research councils [grant NE/N019989/1]. Work was also supported by the National Institute for Health Research Health Protection Research Unit (NIHR HPRU) in Healthcare Associated Infections and Antimicrobial Resistance at the University of Oxford (NIHR200915) in partnership with Public Health England (PHE), and by the NIHR Oxford Biomedical Research Centre (BRC). Computation used the Oxford Biomedical Research Computing (BMRC) facility, a joint development between the Wellcome Centre for Human Genetics and the Big Data Institute supported by Health Data Research UK and the NIHR Oxford Biomedical Research Centre. The computational aspects of this research were funded from the NIHR Oxford BRC with additional support from a Wellcome Trust Core Award Grant [grant 203141/Z/16/Z]. The views expressed are those of the authors and not necessarily those of the NHS, the NIHR, the Department of Health or Public Health England. KCC is Medical Research Foundation-funded. DWC, TEAP and ASW are NIHR Senior Investigators.

## Notes

### Competing Interest Statement

The authors have declared no competing interest.

### Summary of Updates

Original version

