## Supplementary Material for "Niche and local geography shape the pangenome of wastewater- and livestock-associated *Enterobacteriaceae*"

#### Supplementary Tables

Table S1. Metadata for *n*=828 isolates and associated assemblies.
Table S2. Metadata for *n*=2,293 complete plasmids.
Table S3. Information on gene group localization (plasmid/chromosome) for each genus.
Table S4. Antimicrobial resistance (AMR) gene locations within assemblies, as found using abricate.
Table S5. Insertion sequence (IS) locations within assemblies, as found using abricate.

### Supplementary Figures

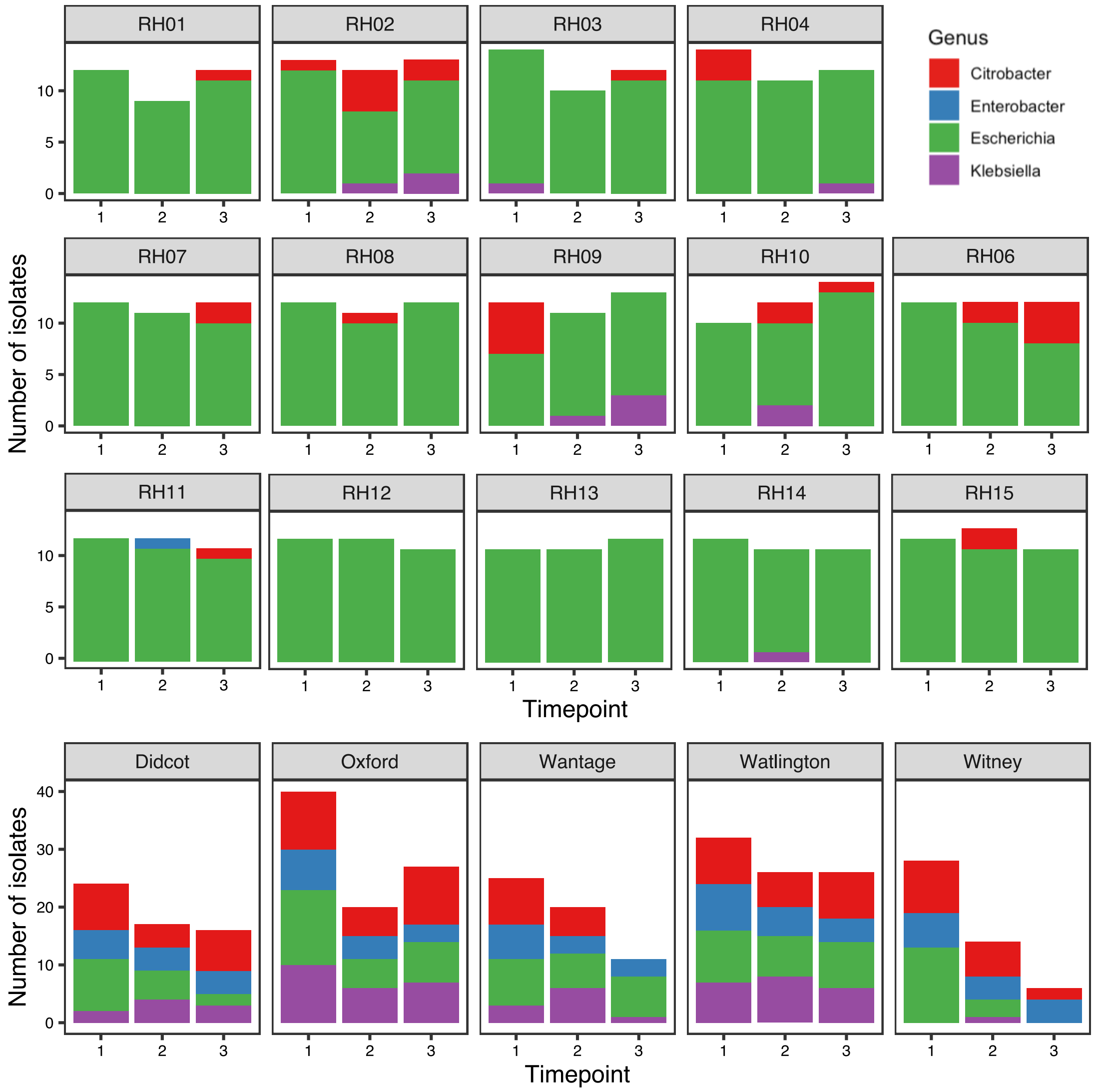


Figure S1. Overview of $n$=828 isolates by sampling location and timepoint. Bars are coloured by genera.

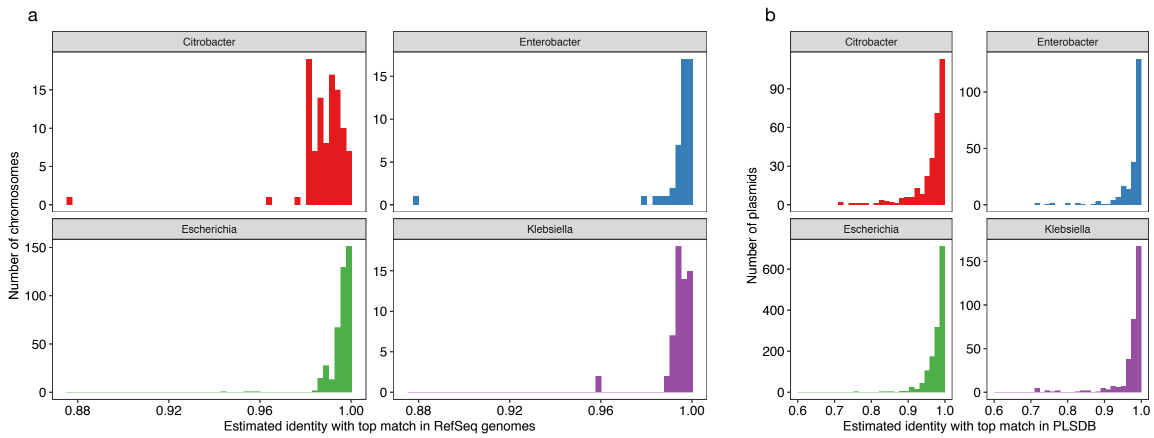


Figure S2. (a) Identity scores (using mash screen) for n=616 circularised chromosomes and their top hit in RefSeq release 88. (b) Identity scores (using mash screen) for n=2,293 plasmids and their top hit in PLSDB v2020_03_04 (20,668 plasmids). Not shown are n=17 plasmids with no match in PLSDB, which were all untyped small plasmids predicted to be non-mobilizable (see Methods).

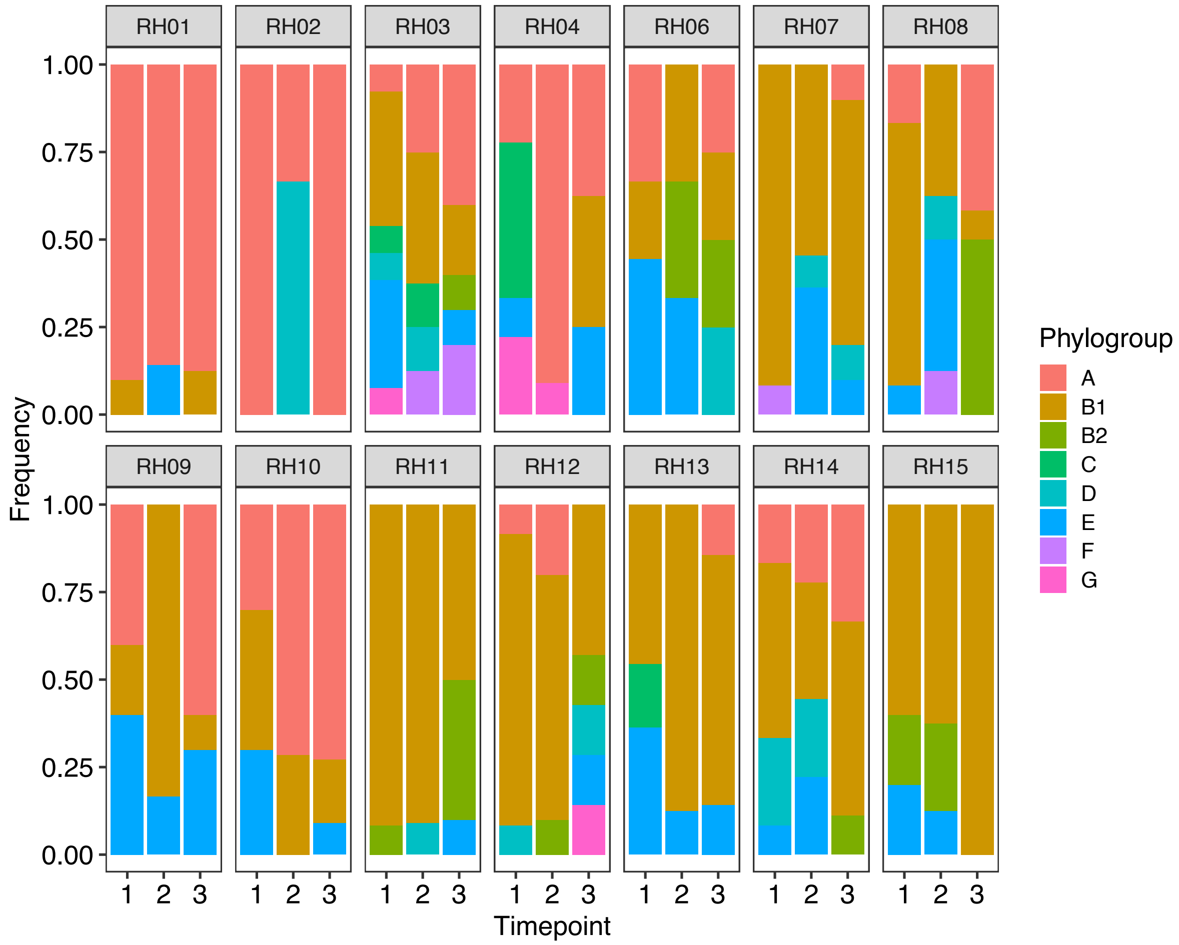


Figure S3. Relative abundance of main *E. coli* phylogroups on livestock farms over time. Proportion of isolates from a timepoint belong to a phylogroup are shown for all *E. coli* isolates within the listed phylogroups (*n*=386). Permutational analysis of variance with adonis, stratifying permutations of farm within livestock niche, showed that livestock niche explained 25.5% of variance (p=0.001) and farm explained 28.1% (p=0.028). Timepoint was not significant (R2=3.6%, p=0.36). Including additional isolates from other or unknown *Escherichia* phylogroups and *E. fergusonii* (*n*=65) did not change the conclusions of the adonis analysis (niche $R^2$=21.1% [p=0.001], farm $R^2$=29.8% [p=0.008], timepoint $R^2$=5.7% [p=0.07]).

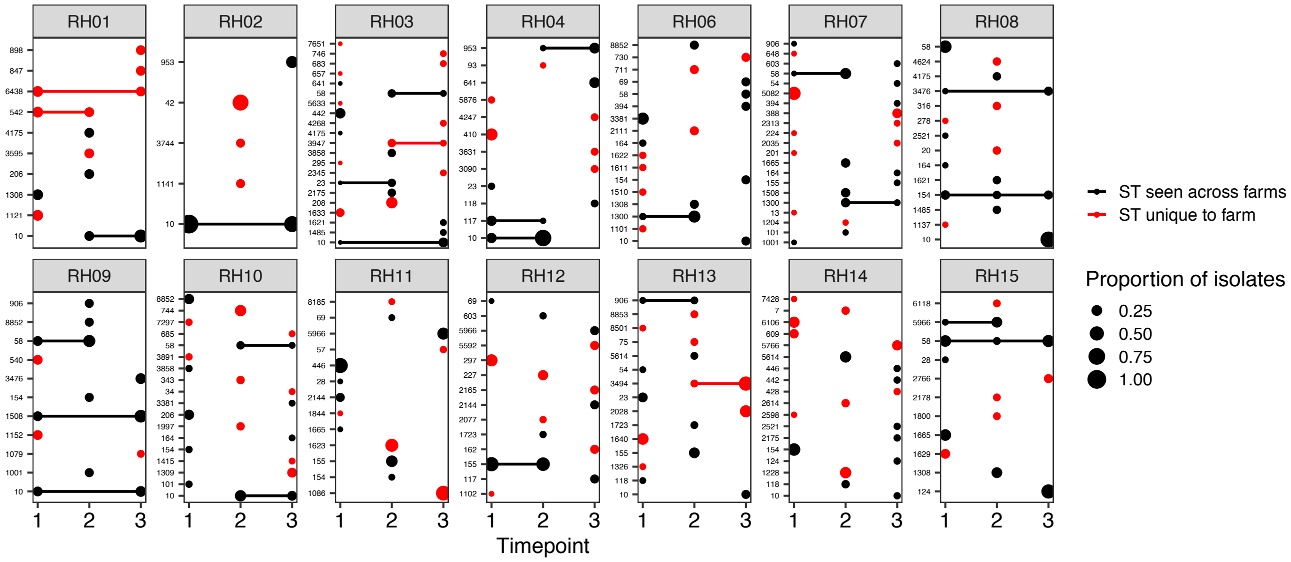


Figure S4. Relative abundance and persistence of *E. coli* STs on livestock farms over time. Size of point shows the proportion of isolates sequenced from that pooled sample. Red points/lines indicate an ST which was only seen in isolates from a single farm; black points/lines indicate STs seen on more than one farm. Only $n$=228 isolates from the main *E. coli* phylogroups (A, B1, B2, C, D, E, F, G) with a known ST are shown, excluding isolates from WwTWs.

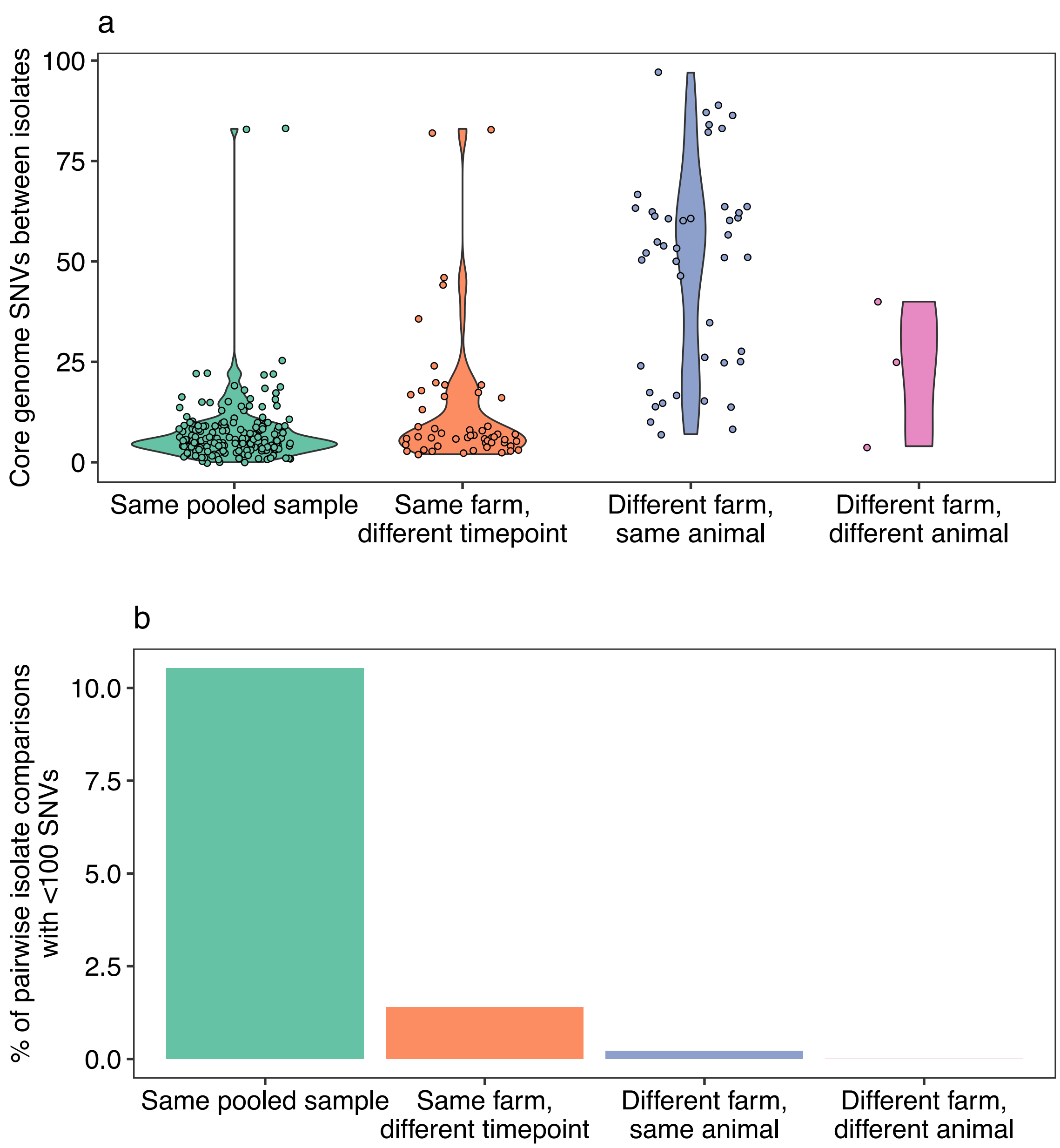


Figure S5. Distributions of pairwise comparisons between *E. coli* isolates from livestock farms. (a) Only comparisons with <100 SNVs are shown. "Same pooled sample" means cultured from the same pooled faecal sample i.e. from the same farm at the same timepoint. Only isolates from farms are shown. (b) Percentage of all possible isolate pairs from a comparison type which had <100 SNVs between them. Isolates from different farms of different animal were very rare (n=3, 0.006%).

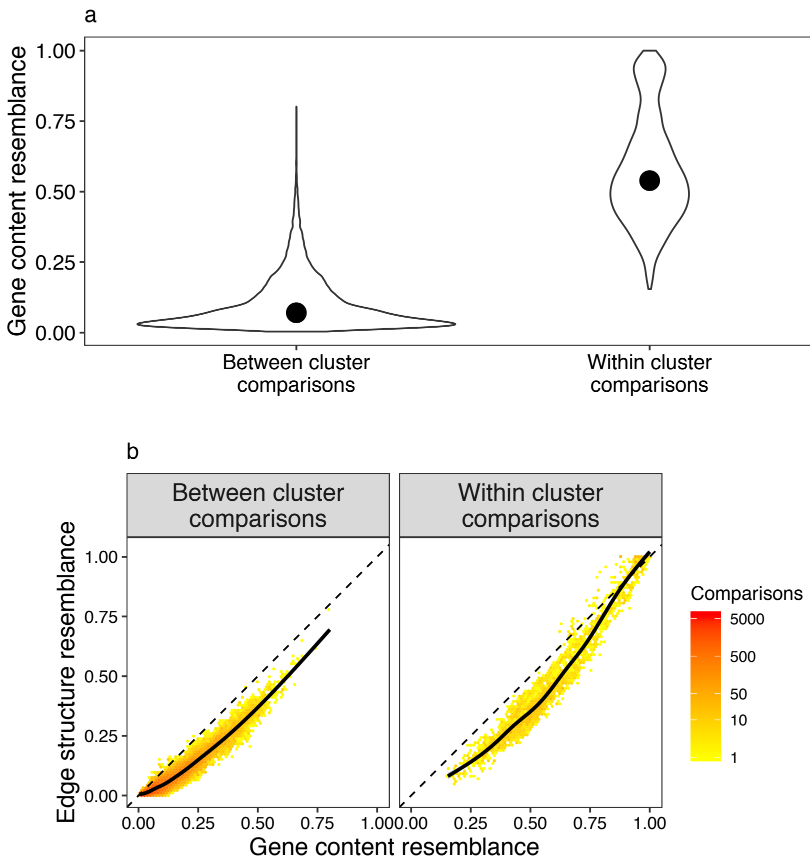


Figure S6. Plasmid clusters defined by distance-free clustering reflect gene content and plasmids show conserved backbones. (a) Comparisons between clusters have a very low median gene content resemblance (median=0.07), compared to within-cluster comparisons (median=0.54) (medians shown by black points). Data is only shown for comparisons between plasmids with 10 or more genes (*n*=848). (b) Gene content resemblance is highly correlated with edge structure resemblance (see Methods). Black line shows a smoothed fit. Colour indicates the number of comparisons contained within a hexagon. Points mainly fall below the line of equality, implying insertions into a plasmid backbone. Where points are above the line of equality, this is due to repeated genetic elements.

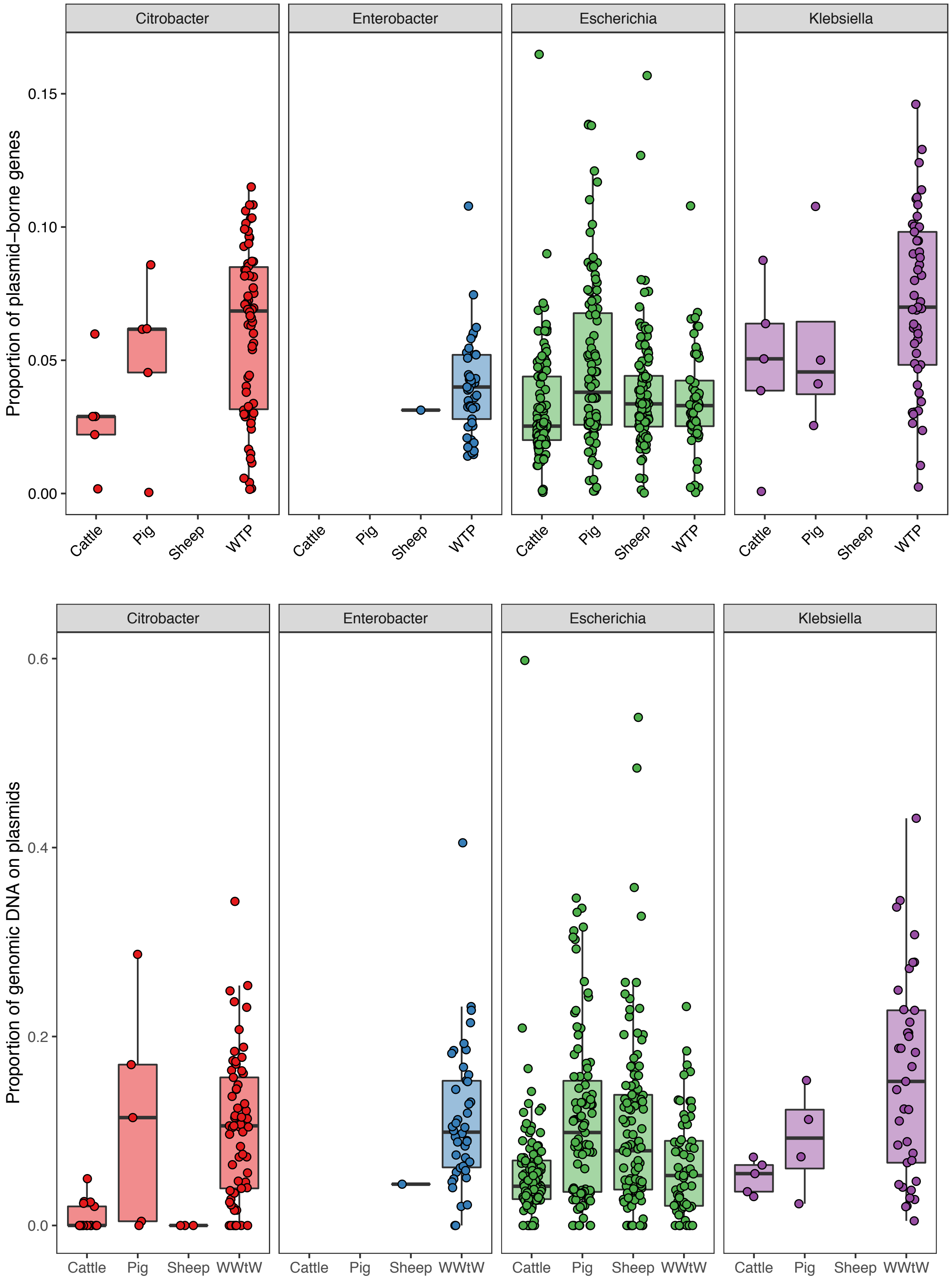


Figure S7. Plasmid-borne proportions of the genome vary between genera and by niche within *Escherichia*. (a) *n*=616 isolates with circularized chromosomes are shown. (b) *n*=558 isolates with complete genomes. Plasmid length was multiplied by each plasmid's normalized depth relative to chromosomal coverage ('copy number') and summed up to give the total proportion of genomic DNA carried on plasmids.

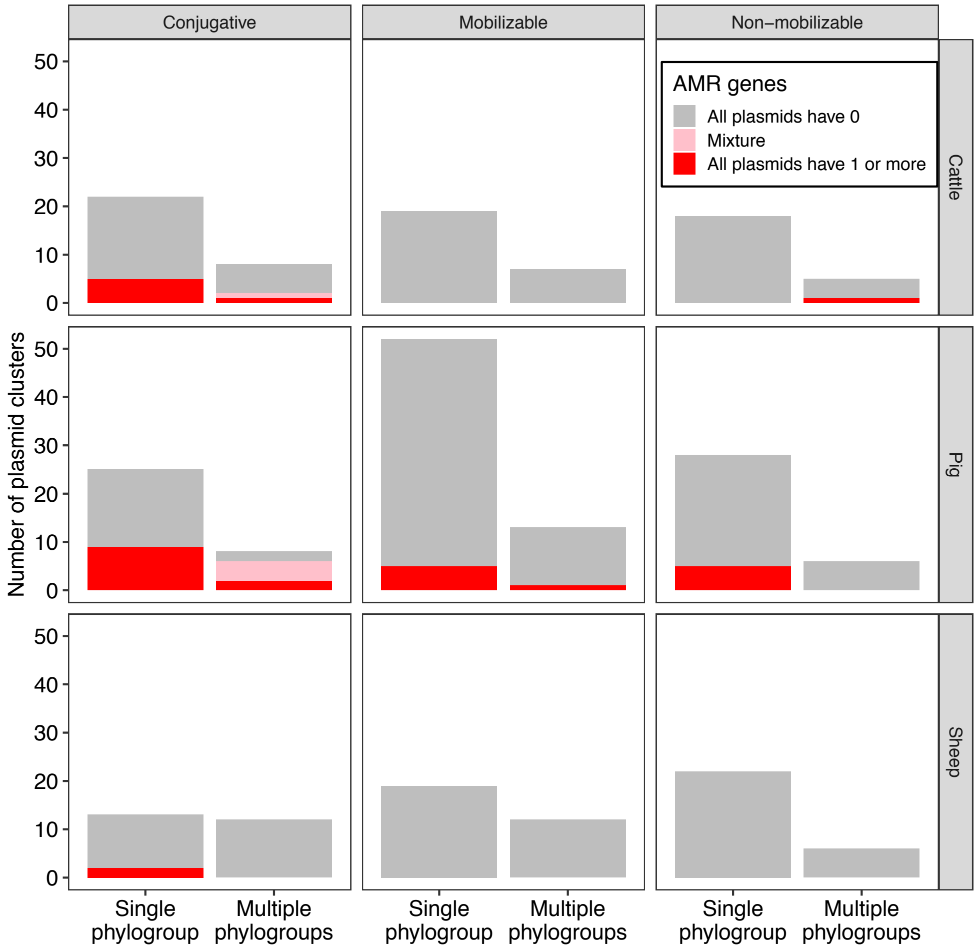


Figure S8. Distribution of 230 plasmid clusters (*n*=1,196 plasmids across 505 isolates) within *Escherichia* on livestock farms. Plasmid clusters were categorised as either appearing only in isolates of a single *Escherichia* phylogroup, or in multiple phylogroups. Plasmid clusters which carried at least 1 AMR gene are shown in red. 220 clusters were consistent for AMR score within a mobility and niche combination (i.e. all plasmids within the cluster had no AMR or AMR), but 10 clusters were heterogeneous (i.e. contained plasmids with no AMR and AMR from the same niche and mobility type: cluster IDs 274, 276, 303, 315, 315, 332, 335, 339, 468, and 564). These were all conjugative plasmids seen in multiple phylogroups, suggesting a broad host range.

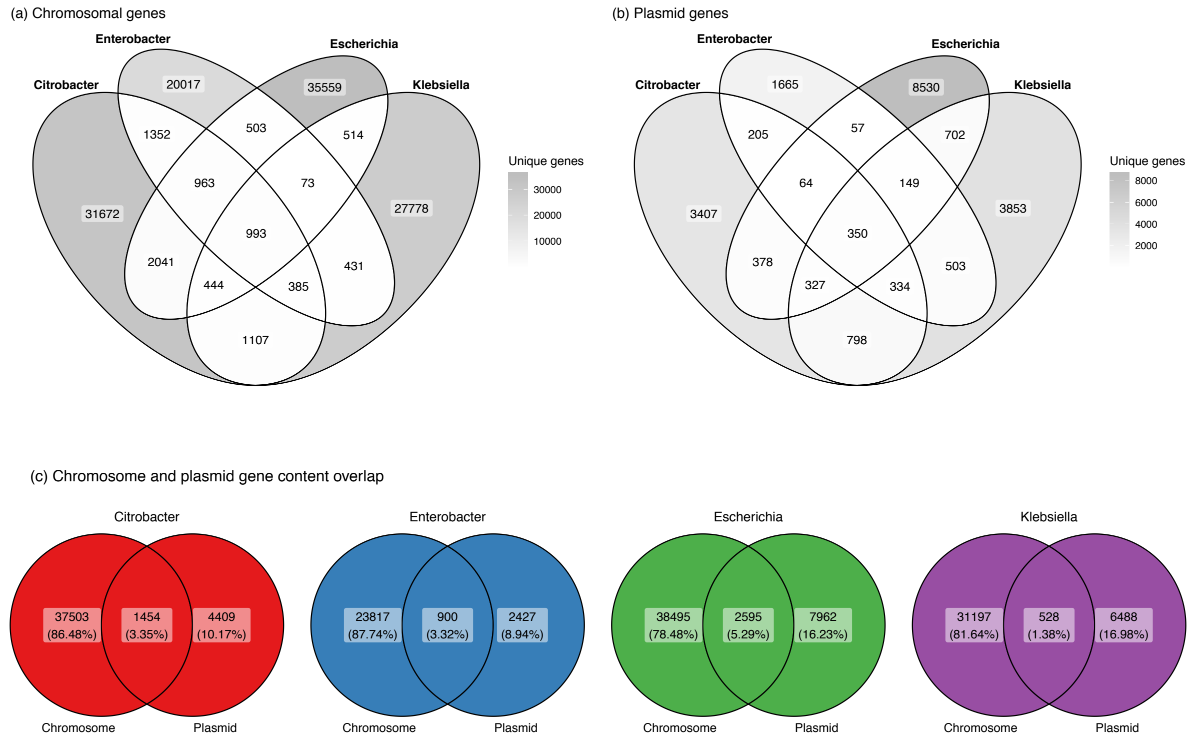


Figure S9. Overlap between genera of chromosomal and plasmid-borne genes. Plasmids carry fewer total genes (Fig. S7) but more diverse and less genus-restricted gene groups. For example, for Escherichia, Genes were clustered into groups at a 95% identity threshold using Roary (see Methods; changing the identity threshold (90%, 99%) did not change the qualitative conclusions) using only isolates with a circularized chromosome (*n*=616), allowing distinguishing between chromosomal and plasmid-borne genes. An overlap is defined as the gene being observed at least once in a given set of samples.

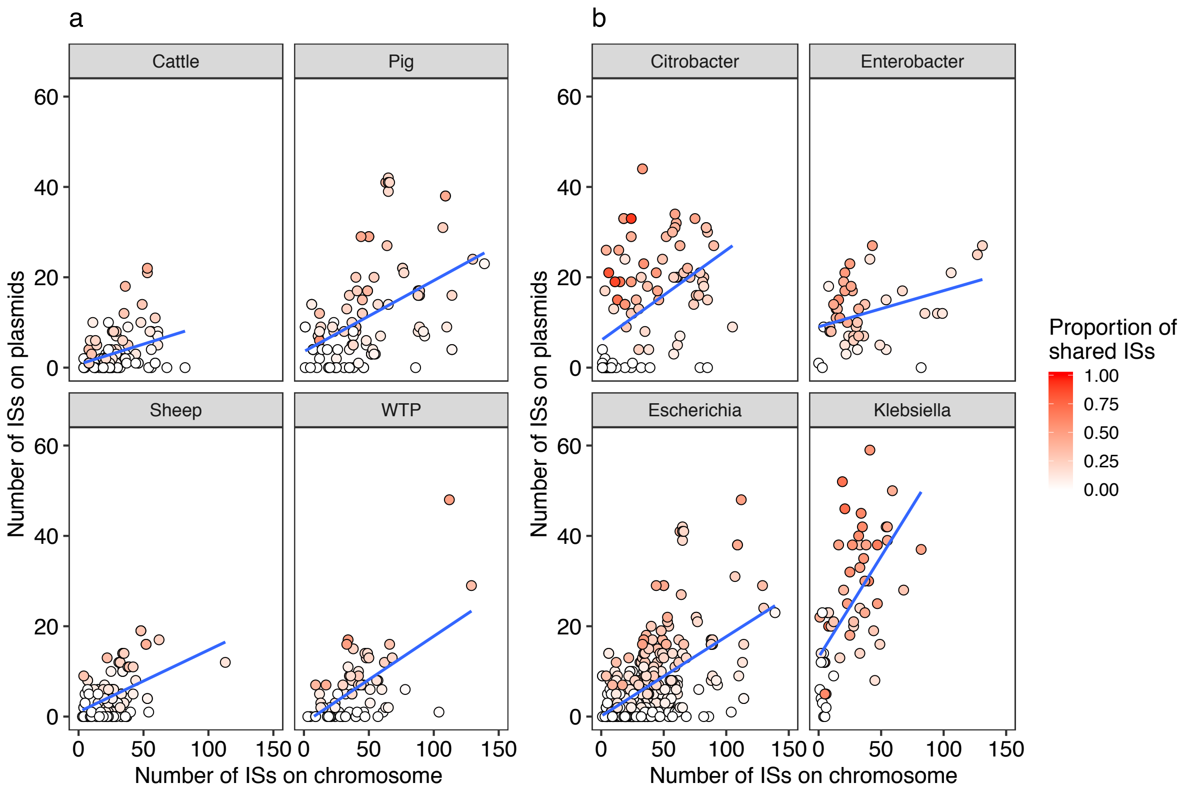


Fig S9. The number of insertion sequences (ISs) on chromosomes is correlated with the number of ISs on plasmids within an isolate. (a) The relationship for *Escherichia* isolates with circularized chromosomes, shown across all four niches. (b) The relationship for all genera. Each point represents one isolate; colours represent the proportion of ISs that are shared across chromosome and plasmid-borne components of an isolate's genome.

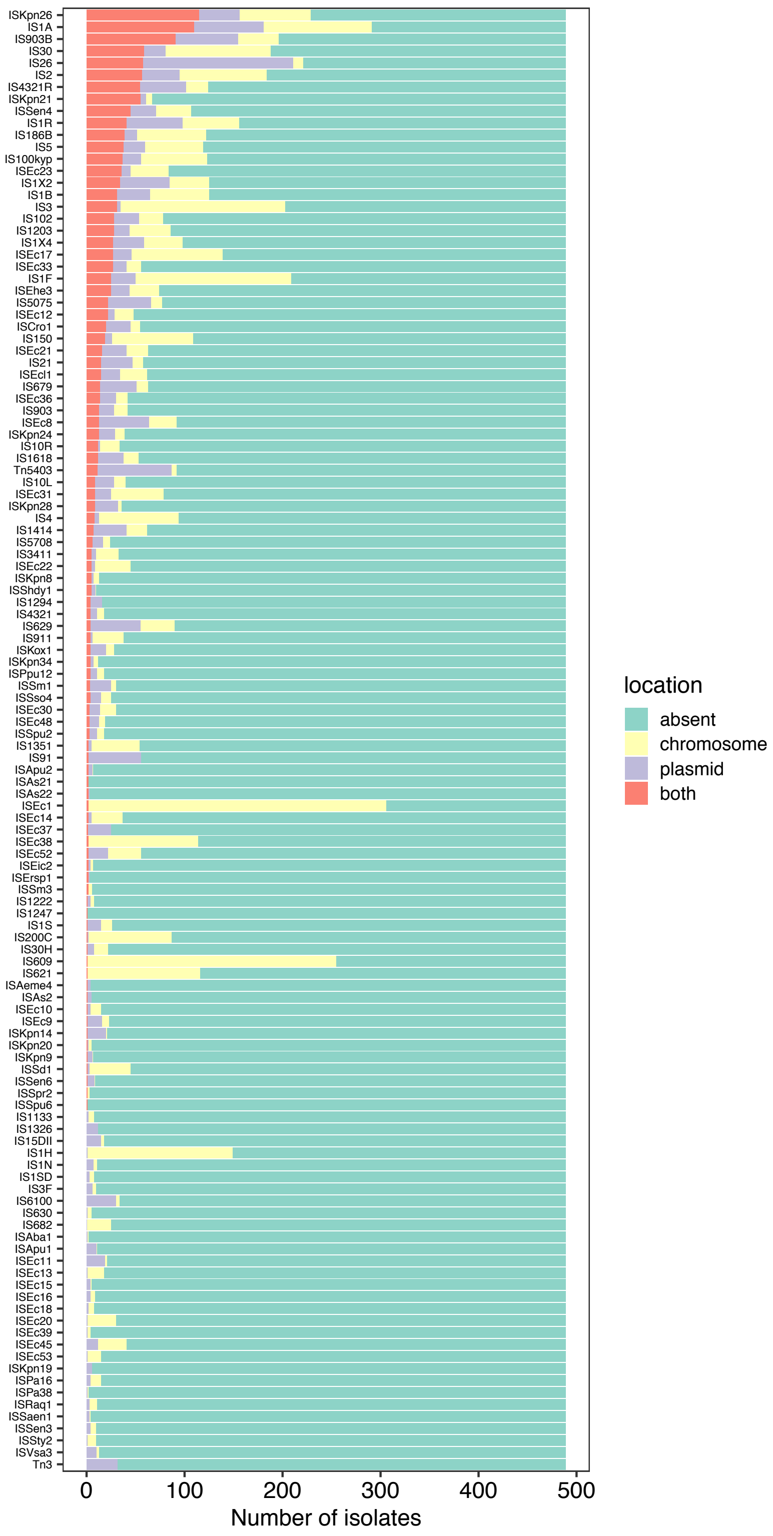


Figure S10. ISs and their locations across isolates of all genera.

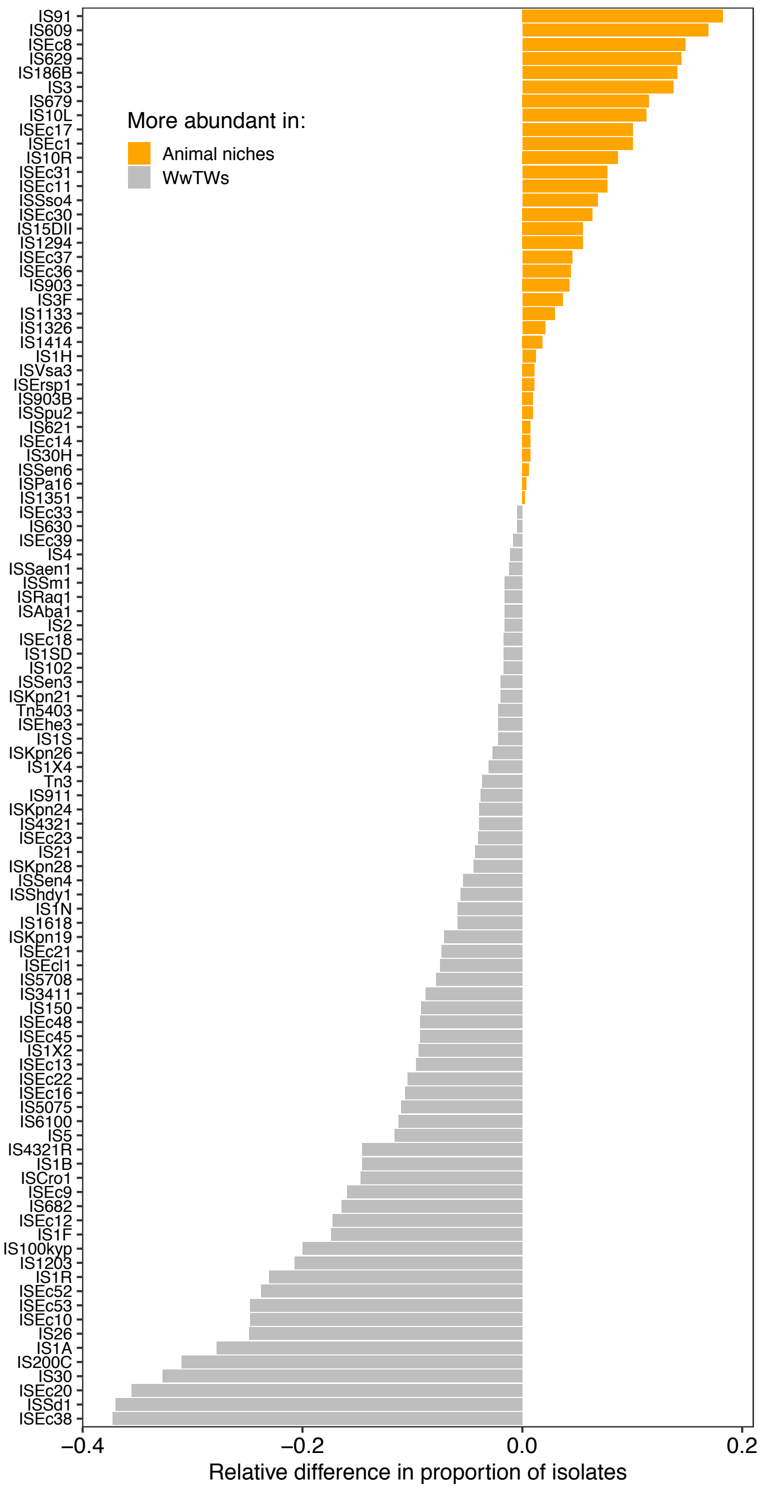


Figure S11. Difference in proportion of *E. coli* isolates containing ISs between WwTWs and other niches (cattle, pig, sheep farms).

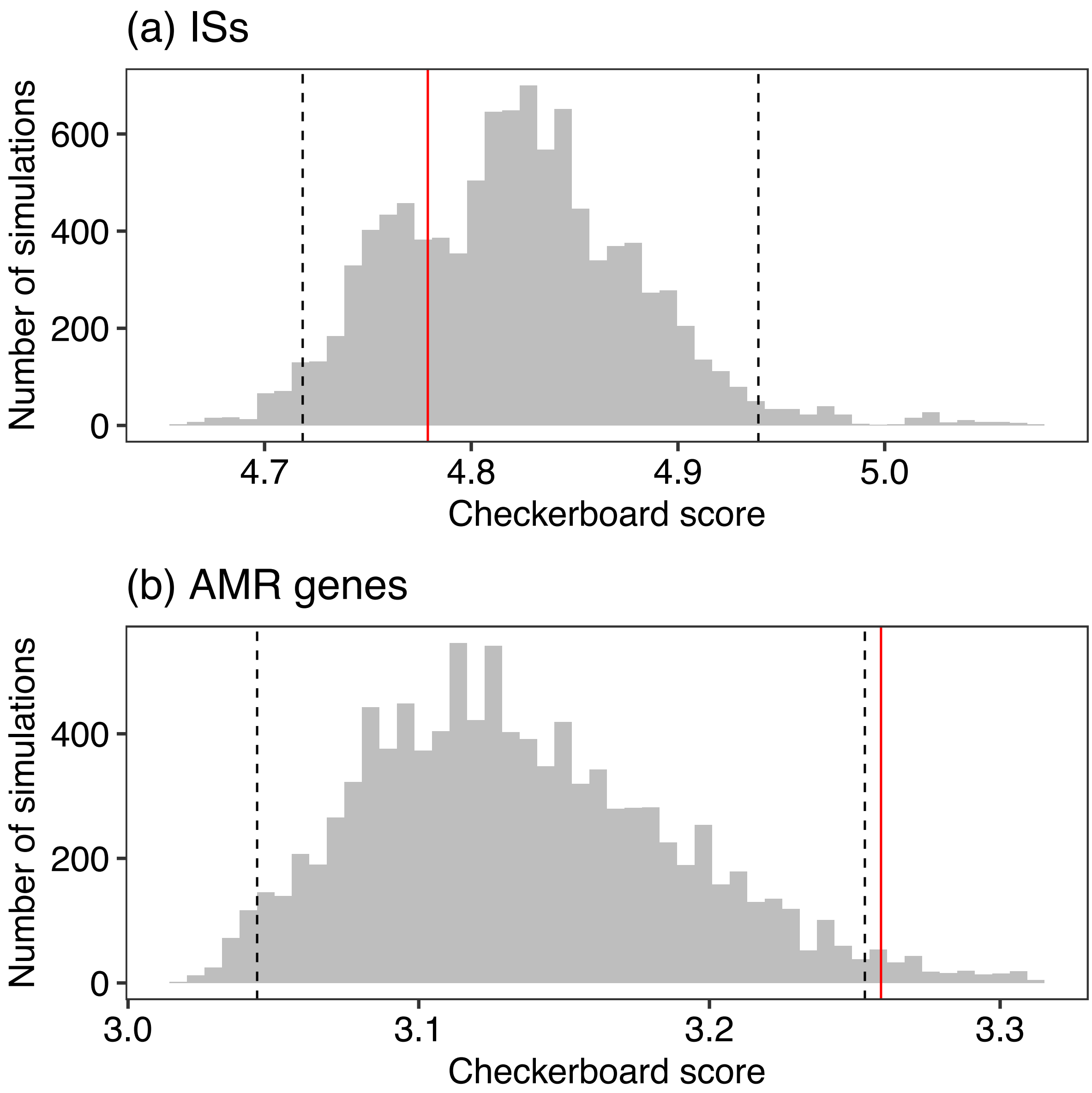


Figure S12. The distribution of (a) ISs and (b) AMR genes across *E. coli* plasmids. Null distribution models for the checkerboard score (grey) for n=10,000 simulations of cooc_null_model in EcoSimR using sim9 method, with dashed lines showed 95% intervals. Red line shows the observed checkerboard score.

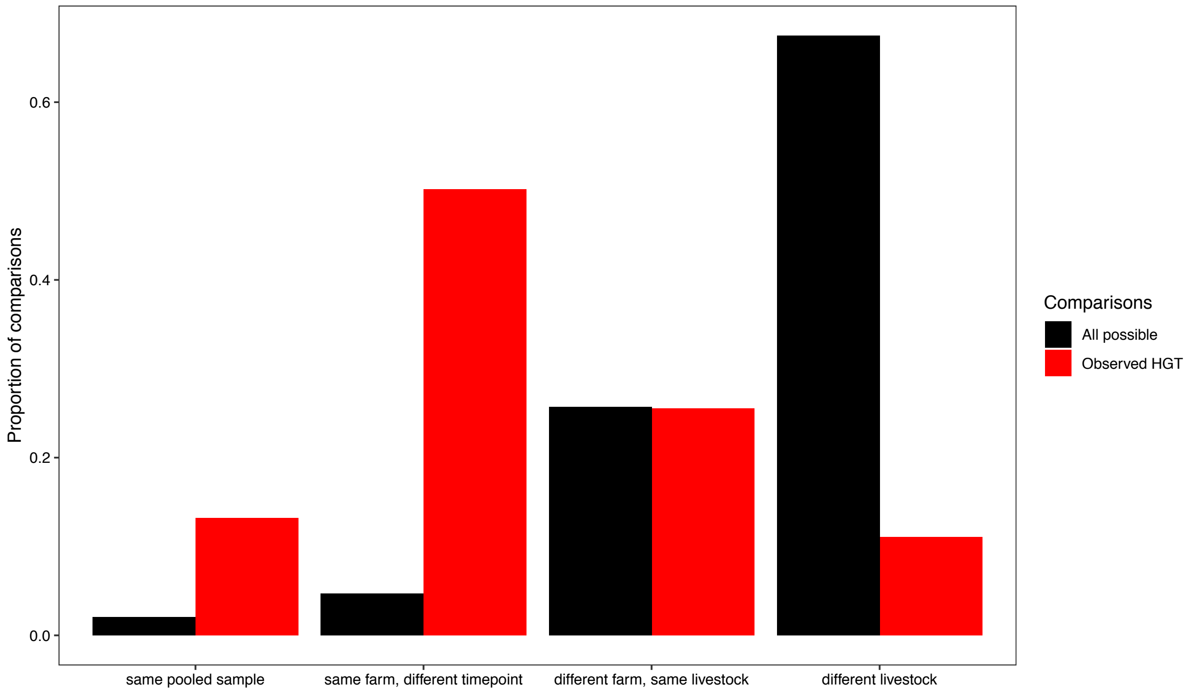


Figure S13. Observed cross-genera HGT event show an overrepresentation of isolates from the same farm, demonstrating the importance of geography for gene movement. Shown are all possible comparisons between farm isolates of different genera (black; *n*=20,723), and only comparisons with at least one possible HGT event of >5,000 bp identical sequence between genomes (red; *n*=235, Table SX). There is an overrepresentation for comparisons from the same farm at the same timepoint (same pooled sample) and at different timepoints, demonstrating a persistent effect of geography.

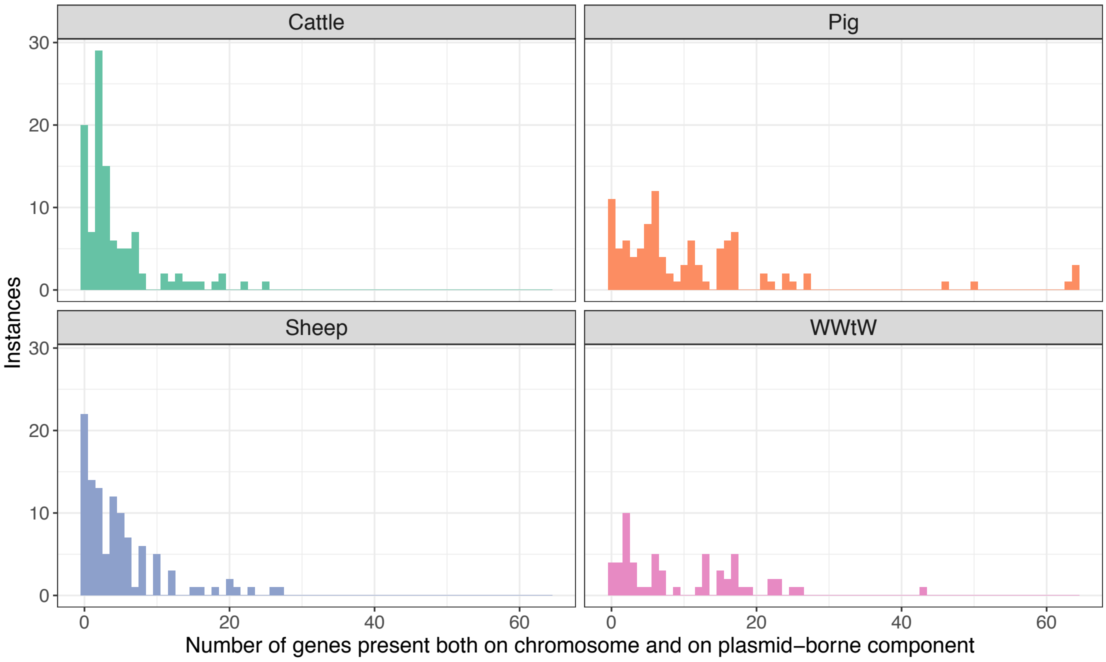


Figure S14. Numbers of genes simultaneously present on both the chromosome and plasmid-borne component within an isolate genome. Data shown for *n*=377 *E. coli* isolates. There are significant differences in the distribution between niches (Kruskal-Wallis test chi-squared=43.4, p<0.001), with pig and WwTW isolates having a more positively-skewed distribution (median 7 and 6.5 respectively, compared to 2 and 3 for cattle and sheep).
